## supplementary information file for "Temporal regularities of vocal exchange in Java sparrows"

### Supplementary Information of "Contagious vocal reaction forms avian vocal exchange: an experimental study in Java sparrows"

Sota Kikuchi, Noriko Kondo, and Hiroki Koda

### Supplementary Section 1. Labelling validations

#### SS.1.1. General

As in the main text, a subset of the data was first extracted: 4 sessions of data were selected from 60 sessions of data to include about 1000 vocalizations, and both evaluators independently labelled the data in the F2F phase. The "agreement" of the labelling between the two labelled data (label-SK, label-NK) was examined.

#### SS. 1.2. Calculating the accuracy rate of the two independent labels

The agreement rate between two annotation labels (label-SK, label-NK) for one segment was defined as the ACCURACY RATE and calculated as follows: $Acc = \frac{N_{agree}}{N_{all}}$, here $Acc$ represents accuracy rate, $N_{all}$ is total number of segmentate vocal region, and $N_{agree}$ is the number of agreements. See results in the main manuscript.

#### SS.1.3. Onset and offset time evaluations

In the two labeled data (label-SK, label-NK), the onset and offset times are recorded, respectively. In order to compare the difference in time, we calculated the time difference ( $\Delta t := t_{sk} - t_{nk}$) from the time in label-SK ($t_{sk}$) to the time in the nearest neighbor label-NK ($t_{nk}$), for onset and offset, respectively. Interval estimation of the obtained time difference data (n=969) was performed using a simple one-sample t-test. If there is a decision bias to annotate label-SK earlier (or later) than label-NK, then there would be a "biased" interval estimation that does not include 0. Otherwise, the interval estimation would include 0.

Figure S.1. shows the distribution of $\Delta t$. The t-test results estimated the interval to be [-0.006, 0.010] and [-0.025, 0.012], for onset time and offset time, respectively, with no statistically significant value bias compared to 0. Accordingly, this result indicates that there was no statistical judgment bias for label-SK and label-NK.

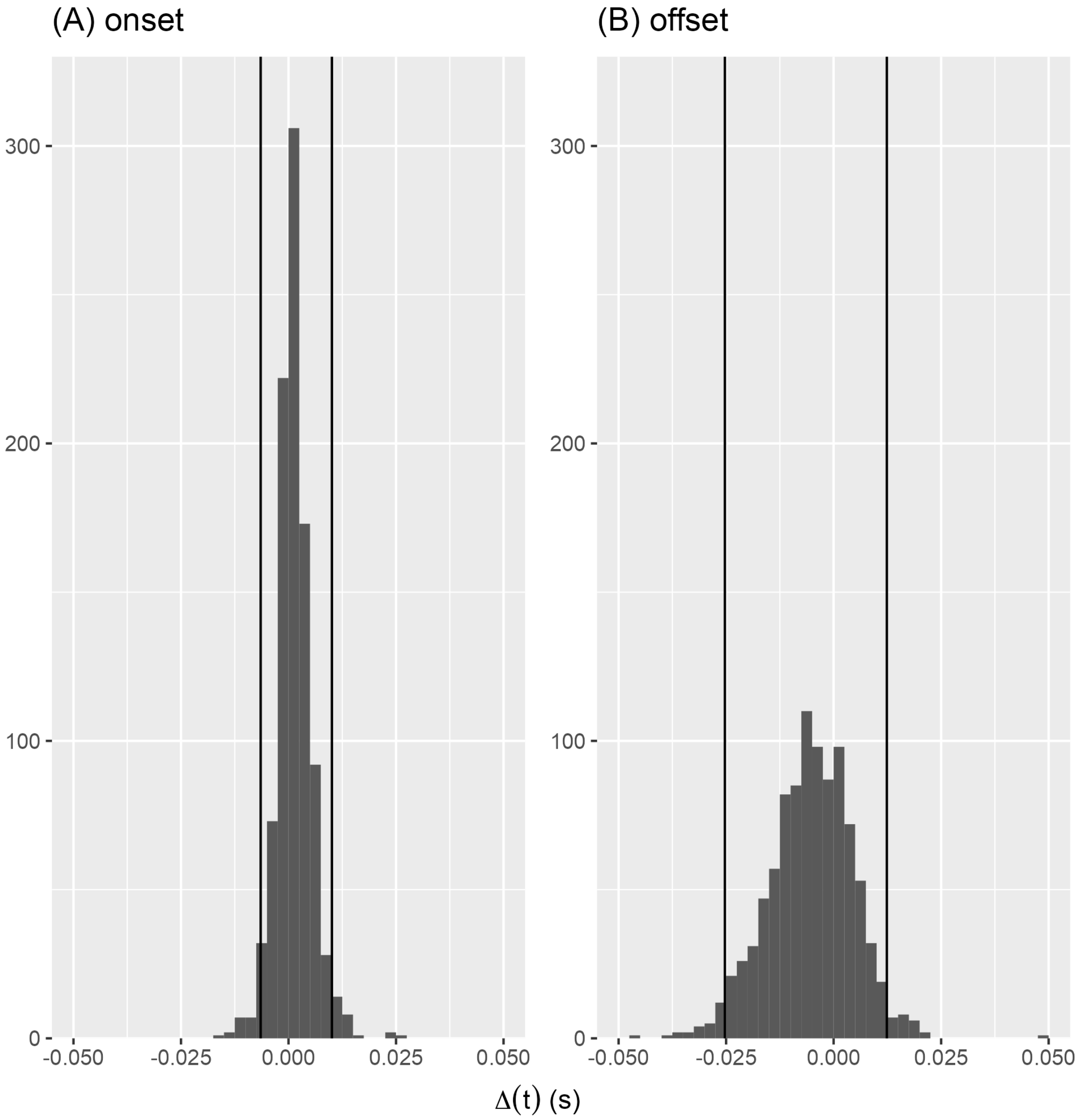


**Figure S.1.** Distributions of the $\Delta t$ of the onset times (A) and offset times (B). The vertical lines represent the lower and upper limits of the 95 percentiles, estimated by the t-test fitted to the obtained $\Delta t$ data.

### Supplementary Section 2. Reports of the time courses of the call rates in the three phases.

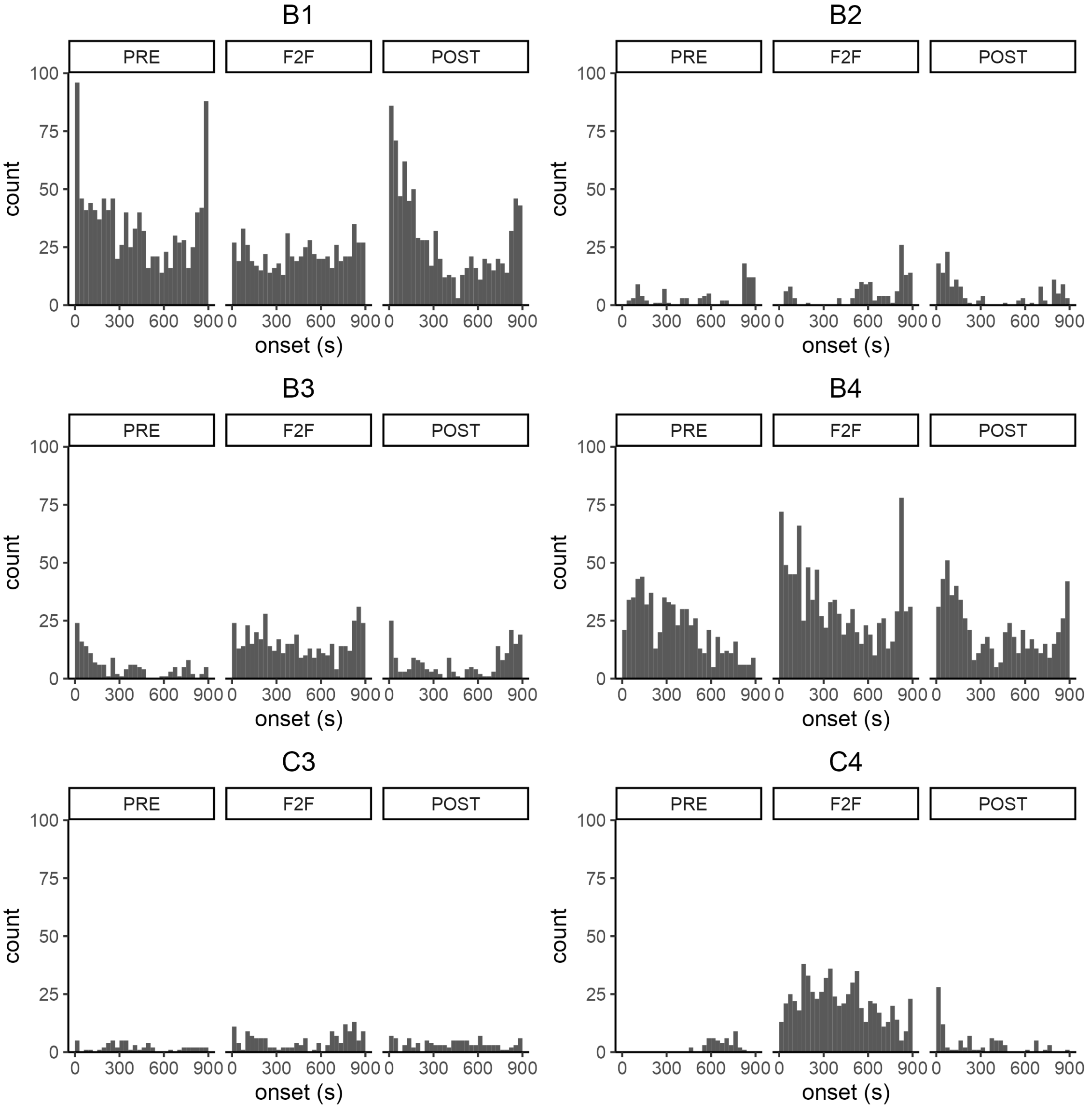


**Figure S2.1.** Time courses of the call rates of the prior individual in the three phases. The time course of the total number of vocalizations for the six individuals (IDs: B1, B2, B3, B4, C3, C4) is shown.

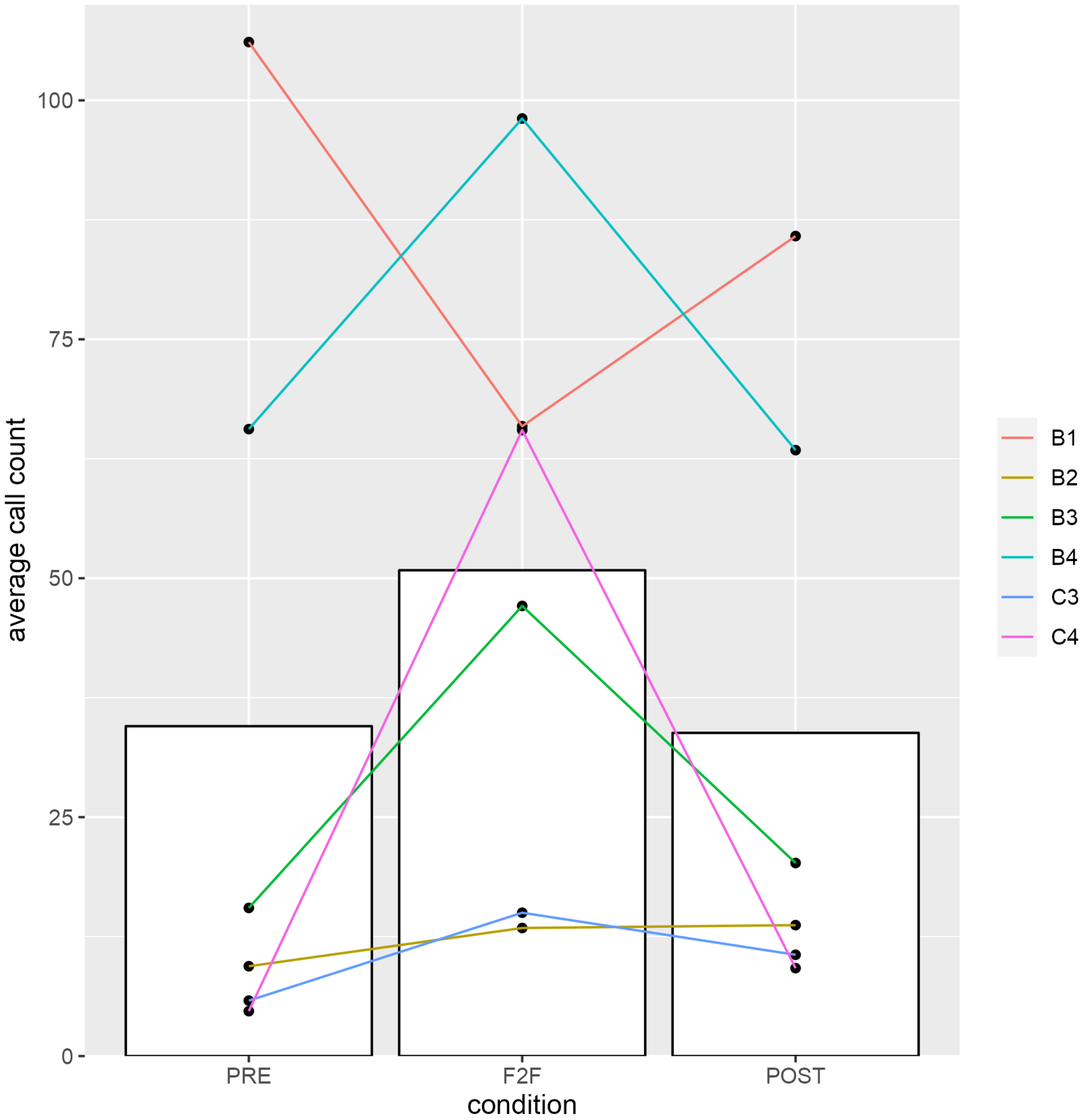


**Figure S2.2.** Plot of the total number of calls of the prior individual per three phases for six individuals.

### Supplementary Section 3. Additional reports for GMM and permutation tests.

#### SS. 3.1. GMM parameter estimations.

Table S3.1. The parameter estimations of the GMMs fitting for the data of exchange transition and repeated transition.

| Data fitted | Data size (N) | Mean (short, long) | Variance (short, long) | Mixture ratio (short, long) |
|---|---|---|---|---|
| Exchange transition | 1710 | -0.728, 0.321 | 0.023, 0.502 | 0.322, 0.678 |
| Repeated transition | 3350 | -0.669, 0.578 | 0.035, 0.296 | 0.167,0.833 |

#### SS. 3.2. Statistical test with permutation tests for the two distributions of exchange and repeated transitions

Figure S3 shows the distribution of the value difference for short components and the 2.5th and 97.5th percentile values generated by the permutation test repeated 10,000 times. The 95th percentile interval includes 0, suggesting that no significant difference was detected between the two distributions of short components.

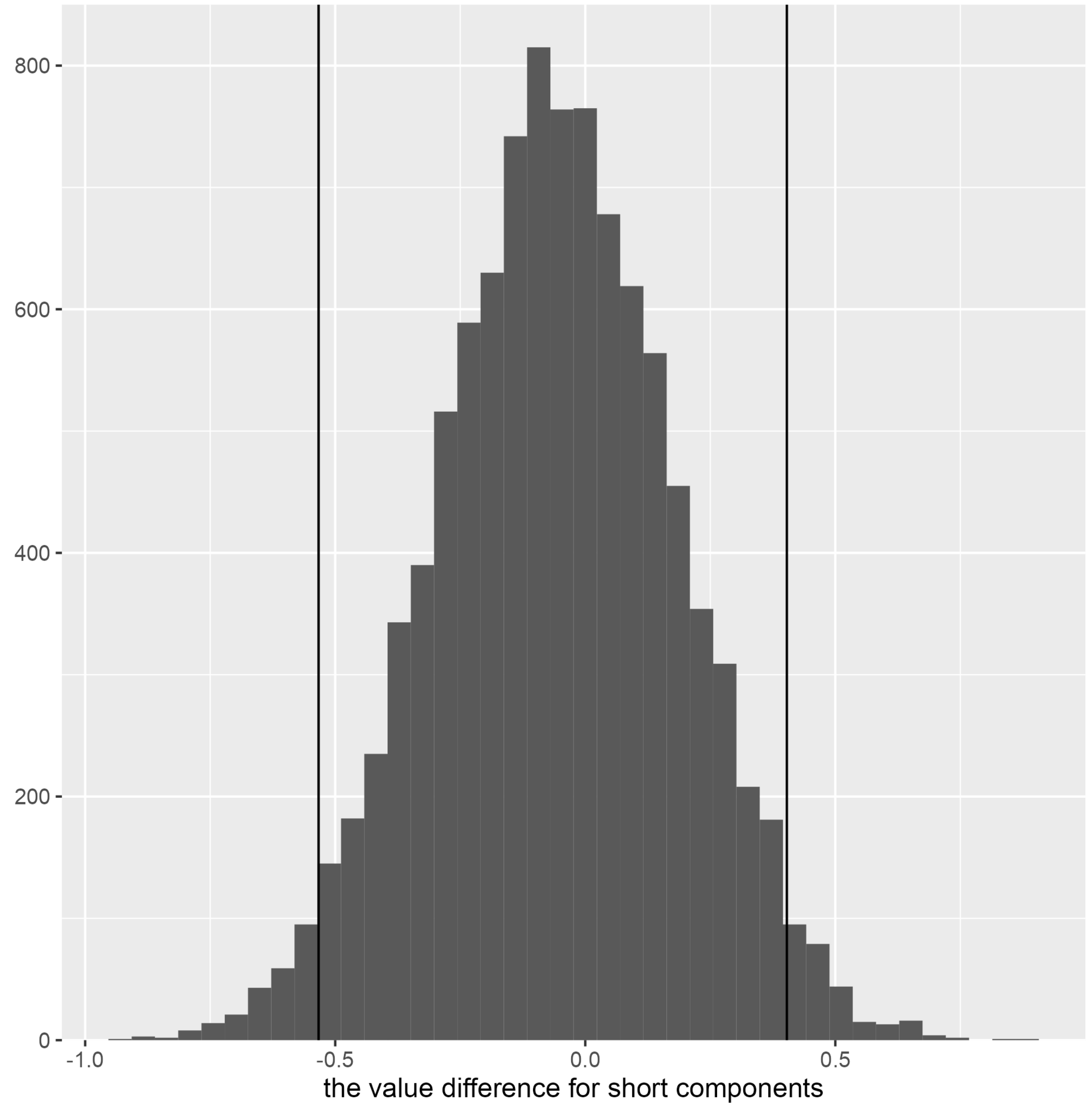


**Figure S3**. The distribution of the value difference for short components and the 2.5$^{th}$ (left side dashed line) and 97.5$^{th}$ (right side dashed line) percentile values generated by the permutation test repeated 10,000 times.
